# Temporal inhibition of electron transport chain attenuates stress-induced cellular senescence by prolonged disturbance of proteostasis in human fibroblasts

**DOI:** 10.1101/2022.11.07.515395

**Authors:** Yasuhiro Takenaka, Ikuo Inoue, Masataka Hirasaki, Masaaki Ikeda, Yoshihiko Kakinuma

## Abstract

We previously developed a stress-induced premature senescence (SIPS) model in which normal human fibroblast MRC-5 cells were treated with either the proteasome inhibitor MG132 or the V-ATPase inhibitor bafilomycin A1 (BAFA1). To elucidate the involvement of mitochondrial function in our SIPS model, we treated cells with an inhibitor of electron transport chain (ETC) complexes I, III, or a mitochondrial uncoupler reagent along with MG132 or BAFA1 and evaluated the induction of premature senescence. SIPS induced by MG132 or BAFA1 was partially attenuated by co-treatment with antimycin A (AA) and rotenone, but not carbonyl cyanide 3-chlorophenylhydrazone (CCCP), in which intracellular reactive oxygen species (ROS) levels, acute mitochondrial unfolded protein responses, and accumulation of protein aggregates were remarkably suppressed. Co-treatment with AA also reversed the temporal depletion of SOD2 in the mitochondrial fraction on day 1 of MG132 treatment. Furthermore, co-treatment with AA suppressed the induction of mitophagy in MG132-treated cells and enhanced mitochondrial biogenesis. These findings provide evidence that the temporal inhibition of mitochondrial respiration exerts protective effects against the progression of premature senescence caused by impaired proteostasis.

**Graphical abstract:** 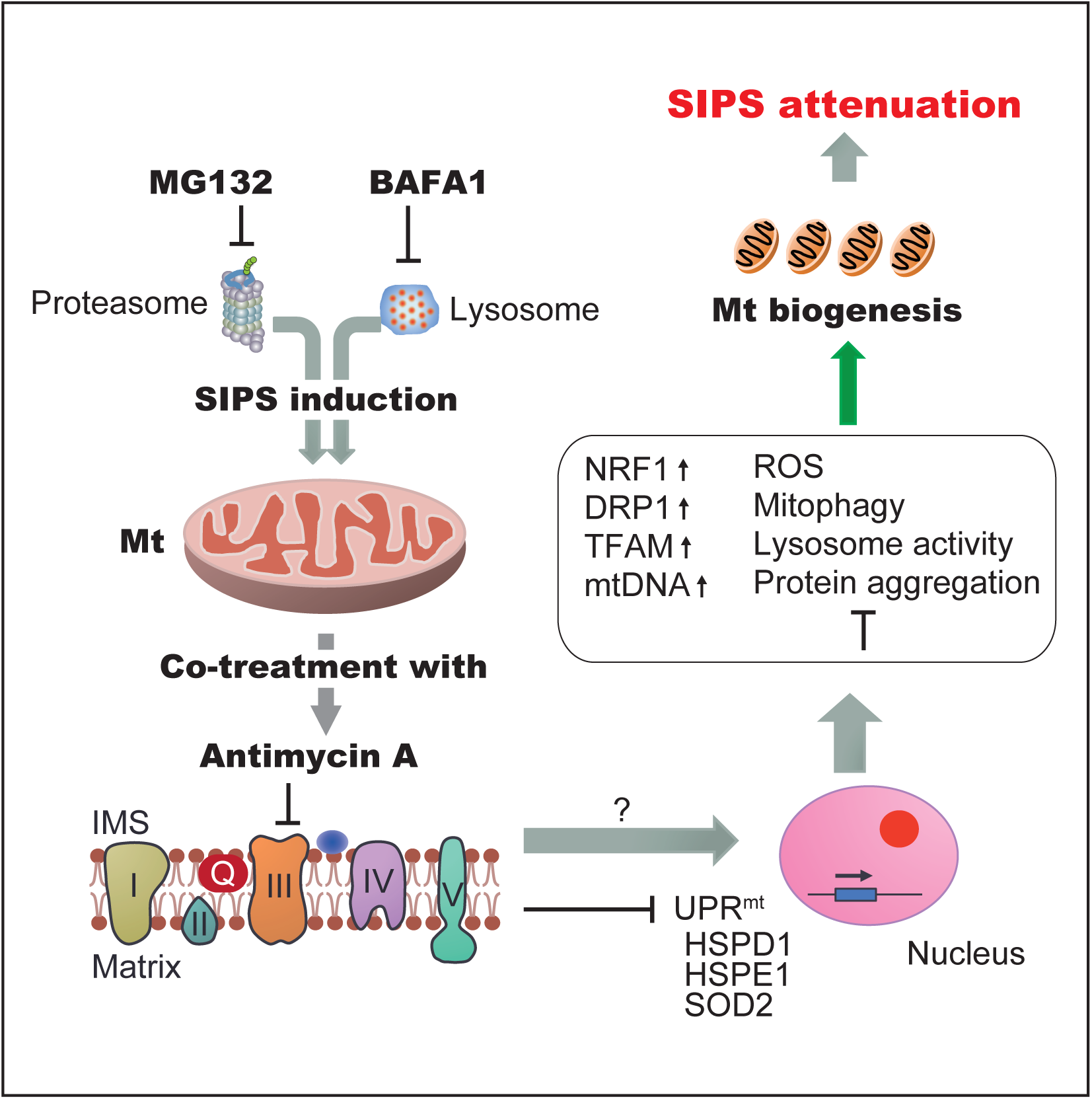
GRAPHICAL ABSTRACT. Cellular senescence is induced by prolonged inhibition of proteasome or lysosome function using MG132 and bafilomycin A1 (BAFA1), respectively. When cells were co-treated with a mitochondrial respiration inhibitor, antimycin A, and MG132 or BAFA1, oxidative stress, mitochondrial unfolded protein response (UPRmt), accumulation of protein aggregates, and mitophagy were suppressed whereas mitochondrial biogenesis was enhanced, resulting in the attenuation of stress-induced cellular senescence (SIPS).

## Introduction

Protein homeostasis, or proteostasis, is a dynamic process that maintains an optimal cellular proteome by modulating protein synthesis and clearance pathways. A decline in proteostasis is a common feature of aging fungi [1], flies [2], nematodes [3], and mammals [4–6]. Reduced proteasome activity results in accumulation of misfolded or damaged protein aggregates in aged organisms. Protein aggregates are also associated with the pathological aspects of neurodegenerative diseases, such as Alzheimer’s and Parkinson’s diseases.

Mitochondria are organelles responsible for the generation of cellular energy by ATP synthesis through the electron transport chain (ETC) in eukaryotic cells and are the main source of reactive oxygen species (ROS) generated in cells [7]. Although mitochondria play a vital role in intracellular energy supply, inhibition of mitochondrial function, especially disruption of the ETC, alternatively leads to lifespan extension in various organisms [8–11]. However, the role of the mitochondria in aging is complicated. Inhibition or disruption of mitochondrial respiratory complexes is known to cause increased levels of ROS [12] and the mitochondrial unfolded protein response (UPR^mt^) [13], which are believed to accelerate aging. In many cases, mitochondria accumulate in senescent cells [14] and show decreased membrane potential [15], which induces mitochondrial dysfunction and increases ROS production [16]. Therefore, a comprehensive understanding of the roles of mitochondria in the process of organismal aging and cellular senescence is required for future aging studies.

Disturbed proteostasis induces stress-induced premature senescence (SIPS) in human fibroblasts [17]. When cells were treated with the proteasome inhibitor MG132, a reversible inhibitor of β1 caspase-like, β2 trypsin-like, and β5 chymotrypsin-like activities of the 20S proteasome, a senescence-like phenotype was induced. We previously established a SIPS model by impairing proteostasis, in which normal human fibroblast MRC-5 cells were treated with a low dose of either MG132 or bafilomycin A1 (BAFA1), a specific and potent inhibitor of vacuolar-type ATPases (V-ATPases) that acidify lysosomes, for five consecutive days [18]. In our SIPS model, the inhibition of mitochondrial function by co-treatment with rapamycin resulted in the recovery of proliferative potential, suppression of excess ROS production, and attenuation of cellular senescence. However, it is unclear whether mitochondrial respiration is directly involved in the SIPS induced by disturbed proteostasis.

In this study, we treated cells with an inhibitor of ETC or a mitochondrial uncoupler reagent along with either MG132 or BAFA1 to reveal the contribution of mitochondrial respiration to SIPS. Among ETC inhibitors, antimycin A (AA), a classical inhibitor of complex III, showed the most prominent effect in preventing SIPS induction. AA also suppressed intracellular ROS levels, acute UPR^mt^ response, accumulation of protein aggregates, and mitophagy, which are commonly observed in MG132-treated cells. Furthermore, AA recovered the temporal depletion of superoxide dismutase 2 (SOD2) in the mitochondrial fraction on day 1 of MG132 treatment and enhanced mitochondrial biogenesis. Our results provide strong evidence that mitochondrial respiration is causally involved in the premature senescence induced by disturbed proteostasis.

## Results

### Antimycin A treatment partially attenuates cellular senescence induced by MG132 or BAFA1

Mitochondrial superoxide levels increase in MG132- and BAFA1-treated MRC-5 cells [18]. To investigate the effect of excess mitochondrial ROS on SIPS induced by MG132 or BAFA1 treatment, we co-treated cells with MitoTEMPO (MT), a mitochondria-targeted superoxide-scavenging reagent, and MG132 or BAFA1 for five consecutive days (Fig. 1A). Although we expected a protective effect of MT against SIPS, the levels of senescence-associated (SA)-β-gal activity were not altered or slightly reinforced by MT co-treatment in a dose-dependent manner (Fig. 1B and 1C). The proliferative activity and senescence-associated morphology at PD5 of MT co-treated cells were also comparable to those of cells treated with MG132 or BAFA1 alone (data not shown). Short-term (1 or 2 days) MT co-treatment also resulted in negative consequences (data not shown). Given the adverse consequences of MT treatment, we investigated the effects of reagents that inhibit the ETC, the main source of mitochondrial superoxide generation. We treated MRC-5 cells with AA and rotenone, which inhibit complexes I and III in the ETC, or the mitochondrial oxidative phosphorylation uncoupler CCCP (1 µM each), respectively, for the first day of five consecutive days of MG132 or BAFA1 treatment (Fig. 1D). Co-treatment with AA and MG132 or BAFA1 remarkably attenuated SIPS induction, whereas the effects of rotenone and CCCP were limited (Fig. 1E, 1F, and 1G), suggesting that inhibiting the function of ETC complex III is the most effective for suppression of SIPS. ETC inhibition by AA was evaluated by measuring the intracellular ATP content. The relative ATP levels of AA co-treated cells were significantly lower than those of MG132-treated cells with no AA at days 1 and 3, presumably because of the prolonged inhibitory effect of AA (Fig. 1H). AA co-treated cells exhibited superior proliferation potential, as assessed by the percentage of Ki-67 positive cells on PD5 (Fig. 1I and 1J). In fact, cells co-treated with AA (MG132+AA) proliferated more vigorously than cells treated solely with MG132 (MG132(-)) after PD5 (Fig. 1K).

**FIGURE 1.**
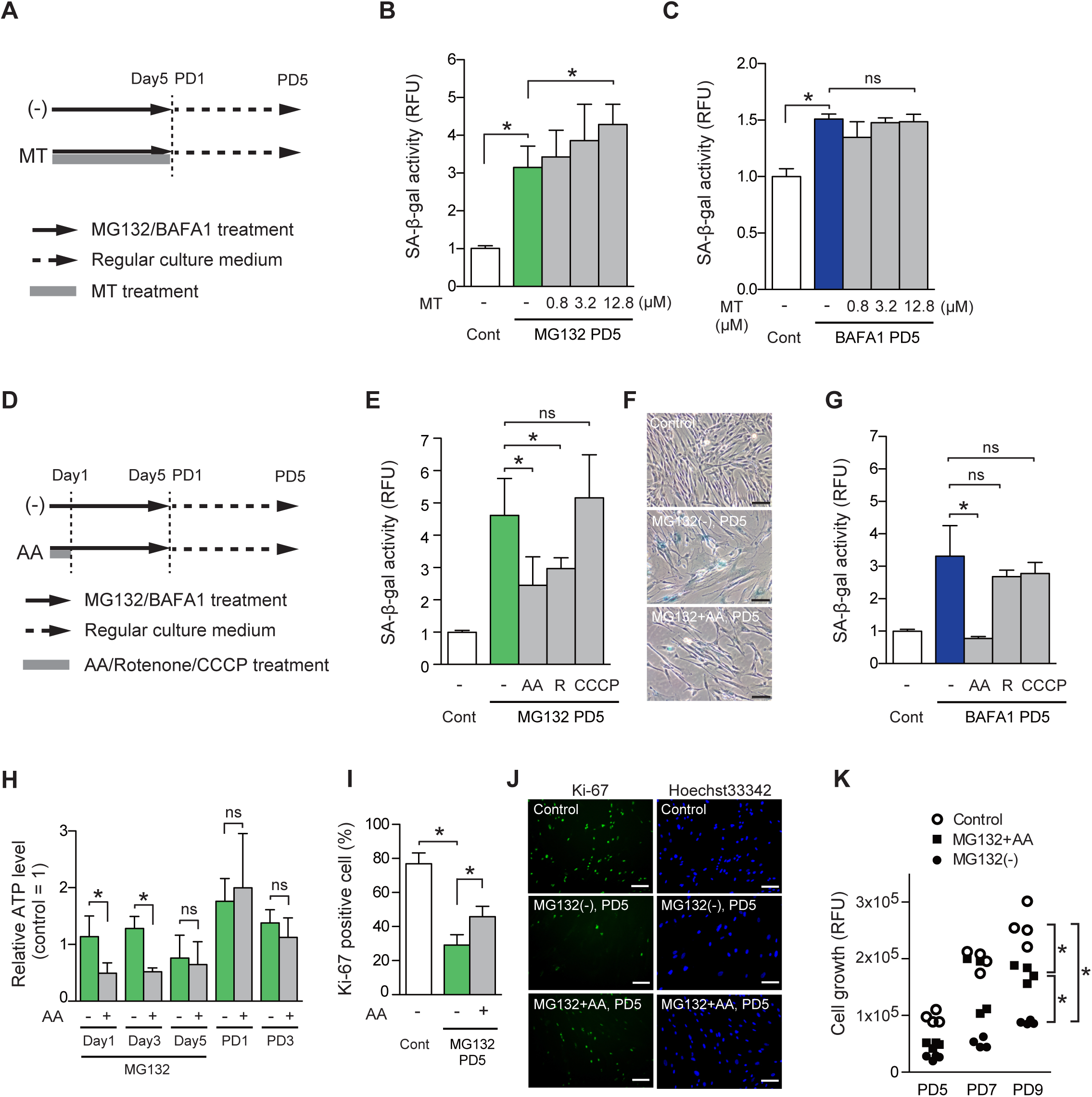
Antimycin A attenuates SIPS induction by prolonged inhibition of proteostasis (A) The experimental design for MitoTEMPO (MT) co-treatment at indicated concentrations (0.8–12.8 µM) with MG132 or BAFA1 in MRC-5 cells. Day, day during drug treatment; PD, post-treatment day. SA-β-gal activity in cells co-treated with MT and MG132 (B) or BAFA1 (C) was measured using the SPiDER-βGal Cellular Senescence Plate Assay Kit. The SPiDER-βGal fluorescence was normalized to that of Hoechst 33342-stained nuclei. Cont, untreated control cells. Values are shown as the mean (SD) (*n* = 4). The asterisk (*) indicates *p* < 0.05, as determined by one-way ANOVA, followed by Tukey’s multiple comparison test. (D) Experimental design for ETC inhibitors, 1 µM antimycin A (AA), rotenone (R), or CCCP, with MG132 or BAFA1 in MRC-5 cells. (E) SA-β-gal activity of cells co-treated with each of the three ETC inhibitors and MG132. (F) Representative images of SA-β-gal-stained MRC-5 cells co-treated with MG132 and AA on PD5. Scale bar = 100 µm. (G) SA-β-gal activity of cells co-treated with each of the three ETC inhibitors and BAFA1. (H) Intracellular ATP content was measured using a luciferase-based assay. The luminescence of ATP was normalized to the cellular DNA concentration that was quantified using SYBR Gold. Values are shown as the mean (SD) (*n* = 4). The asterisk (*) indicates *p* < 0.05 for MG132-treated (-) vs. AA co-treated (+) cells. (I) Percentages of Ki-67 positive cells on PD5. Values are shown as the mean (SD) (*n* = 5). (J) Representative images of Ki-67-stained MRC-5 cells co-treated with MG132 and AA on PD5. Scale bar = 100 µm. (K) Cell proliferation was evaluated by measuring Hoechst 33342 fluorescence from PD5 to PD9. The asterisk (*) on PD9 indicates *p* < 0.05, using one-way ANOVA with Tukey’s multiple comparison test.

### Antimycin A suppresses intracellular ROS production and attenuates oxidative stress responses in MG132-treated cells

Prolonged treatment of MRC-5 cells with MG132 or BAFA1 increases the intracellular ROS levels during and after treatment [18]. It is also known that treatment of cells with AA induces excess ROS production [12]. We first confirmed that a 1-day treatment of AA at 1 µM enhanced the production of mitochondrial superoxide in MRC-5 cells (Fig. 2A). We next performed time-course analyses of mitochondrial superoxide levels in MG132 and AA co-treated cells. Mitochondrial superoxide level increased as early as 12 h after MG132 treatment. By AA co-treatment, the levels were significantly suppressed at 12 h and day 5 (Fig. 2B). Intracellular H_2_O_2_ levels were also significantly suppressed by AA co-treatment on days 1 (Fig. 2C and 2D) and 3 (Fig. S1A and S1B). The protein level of NRF2, a redox-sensitive transcription factor, transiently increased during the early period of MG132 treatment (Fig. 2E and 2F). AA co-treatment suppressed the sudden increase in NRF2 at 3 h, indicating that AA co-treated cells promptly responded to and alleviated the oxidative stress caused by MG132 (Fig. 2E and 2F). Protein levels of heme oxigenase-1 (HO-1), a downstream factor of NRF2, were also suppressed at 6 and 24 h by AA co-treatment. Taken together, the oxidative stress responses caused by MG132 treatment were significantly suppressed by co-treatment with AA.

**FIGURE 2.**
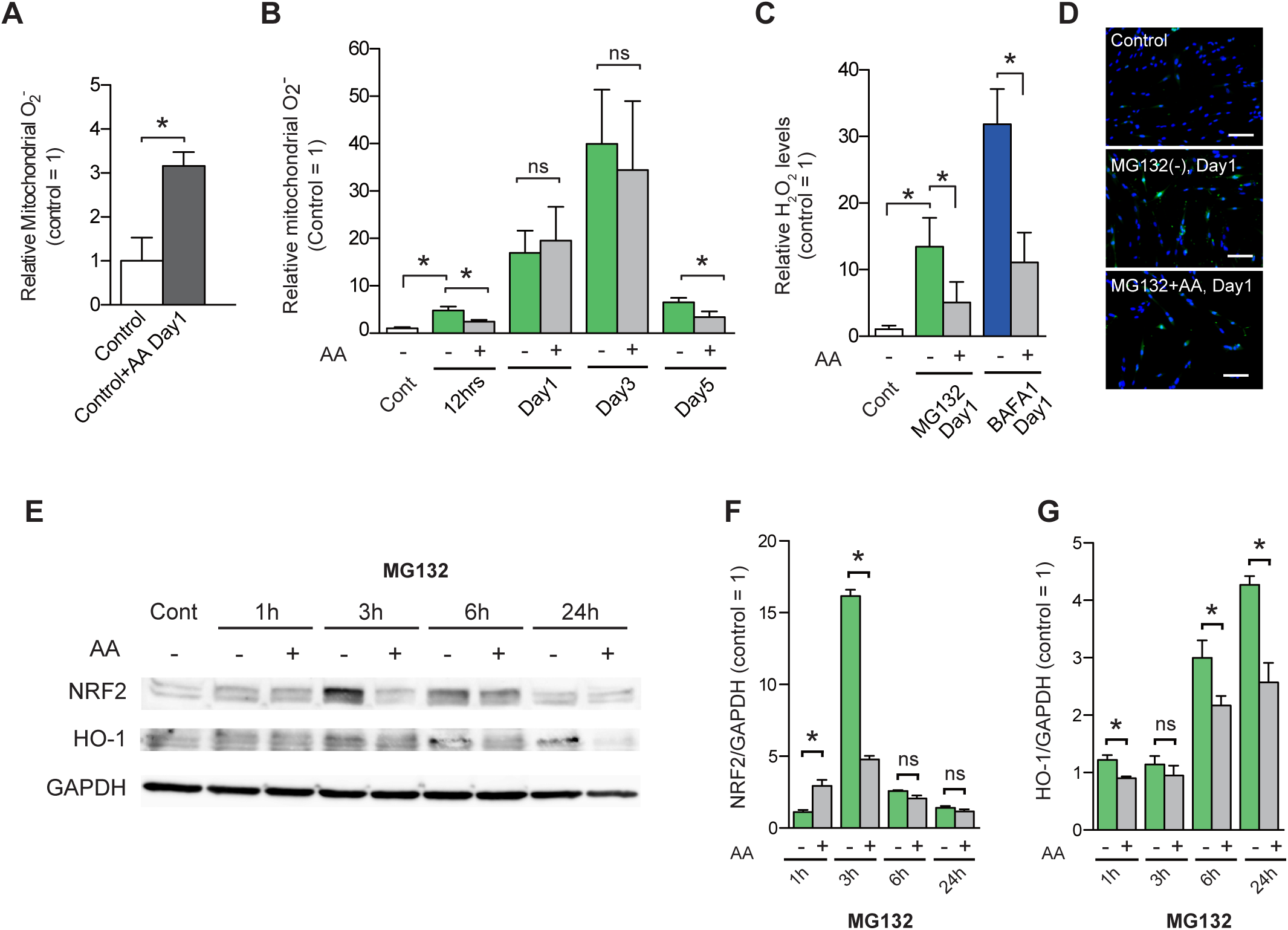
Antimycin A suppresses generation of excess mitochondrial superoxide and acute oxidative stress responses induced by prolonged inhibition of proteostasis (A) Relative mitochondrial superoxide levels were examined using MitoSOX Red mitochondrial superoxide indicator. Cells solely treated with AA induced generation of excess mitochondrial superoxide on day 1. (B) Relative mitochondrial superoxide levels in MG132-treated (-) and AA co-treated (+) cells. The fluorescence intensity for MitoSOX was normalized to the number of nuclei. Relative values are shown as the mean (SD) from the four different images. The asterisk (*) indicates *p* < 0.05 by a Mann-Whitney *U* test. (C) Relative intracellular H_2_O_2_ levels in MG132-treated (-) and AA co-treated (+) cells on day 1. The fluorescence intensity for HYDROP was normalized to the number of nuclei. Relative values are shown as the mean (SD) (*n* = 4). (D) Representative images of control, MG132-treated, and AA co-treated cells on day 1 stained with HYDROP (green) and Hoechst 33342 (blue). Scale bar = 50 µm. (E) Suppression of NRF2 and HO-1, markers for oxidative stress, were examined by the immunoblot analyses of MG132-treated and AA co-treated cells. Cont, untreated control cells. Quantitative analyses of immunoblots for NRF2 (F) and HO-1 (G). Intensities of immunoblot bands were normalized to those of GAPDH, respectively. Relative values are shown as the mean (SD) (*n* = 4).

### Antimycin A suppresses the accumulation of protein aggregation in MG132-treated cells

MG132 markedly increases intracellular protein aggregates during and even after the treatment [18]. AA co-treatment significantly suppressed the accumulation of protein aggregates on day 3 and PD5 (Fig. 3A and 3B). To investigate whether the suppression of protein aggregates was mediated by enhanced protein clearance, we analyzed the levels of lysosomal acidity and LC3-II protein, the active form of LC3. The acidic lysosomal signal significantly increased in MG132-treated cells on day 3, but not in AA co-treated cells (Fig. 3C and 3D). The LC3-II/LC3-I ratio increased in MG132-treated cells on days 3 and 5, but not in AA co-treated cells (Fig. 3E and 3F), suggesting that the reduction in protein aggregates by AA was not caused by enhanced proteolytic processes. The chaperone protein HSP70 was dramatically upregulated as early as 6 h by the treatment with MG132, but was significantly suppressed by AA co-treatment at 6 h, days 3 and 5, compared with MG132-treated cells (Fig. 3E and 3G). p62/SQSTM1 accumulated by MG132 treatment, but the protein levels were not altered by co-treatment with AA (Fig. 3E). Taken together, these results indicate that the reduced levels of intracellular protein aggregates observed in AA co-treated cells were neither mediated through increased autophagic flux nor through increased chaperone protein, but possibly derived from limited generation of aggregated proteins caused by ROS.

**FIGURE 3.**
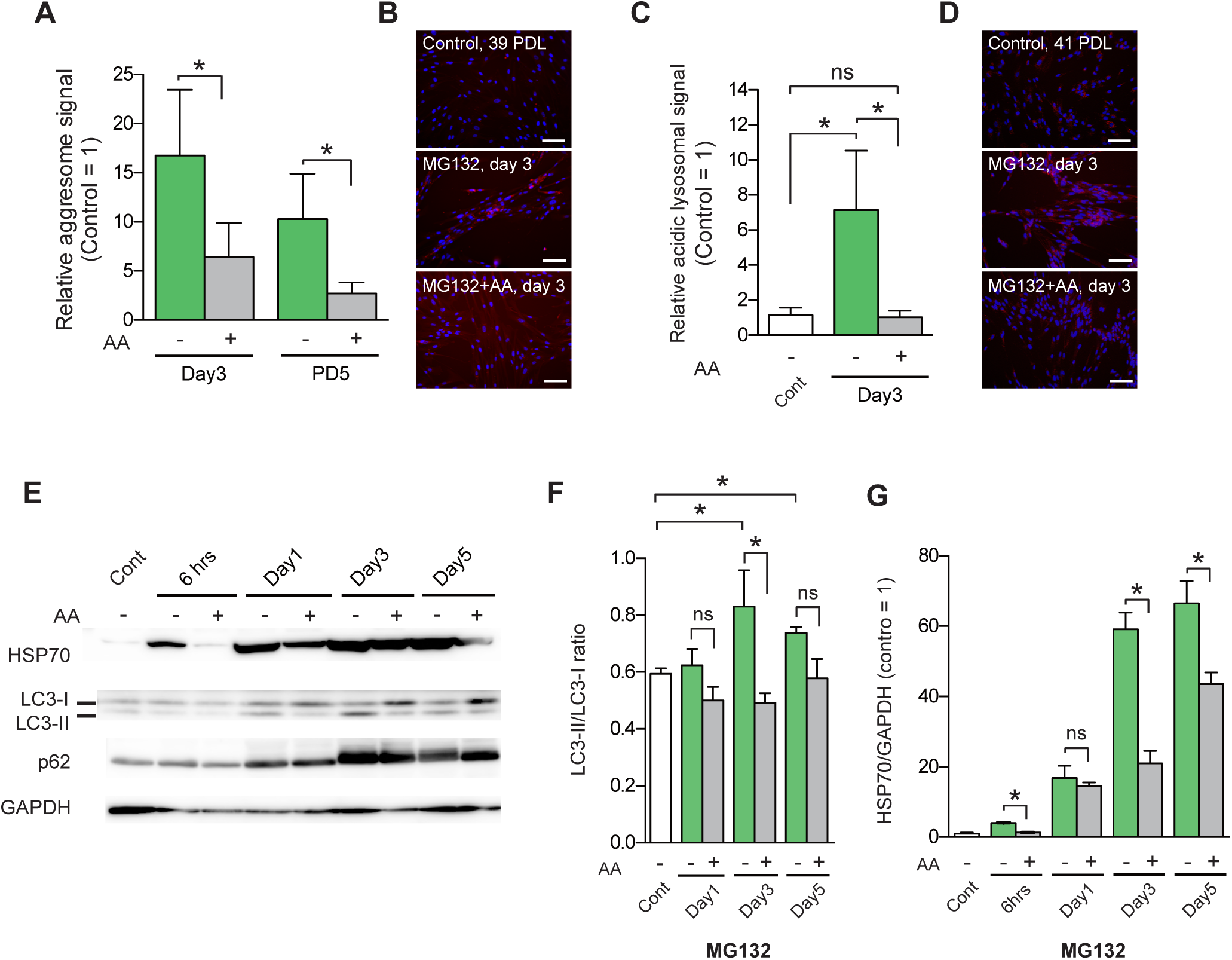
Antimycin A suppresses accumulation of intracellular protein aggregates induced by prolonged inhibition of proteostasis (A) The fluorescence intensity for intracellular protein aggregates was normalized to the number of nuclei. Relative values are shown as the mean (SD) from the measurements of the four different images. The asterisk (*) indicates *p* < 0.05 by a Mann-Whitney *U* test for MG132-treated (-) vs. AA co-treated (+) cells. (B) Representative images of accumulated protein aggregates in controls (39 PDL), MG132-treated, and AA co-treated cells on day 3. Cells were stained with ProteoStat Aggresome Detection Reagent (red) and Hoechst 33342 (blue). Scale bar = 50 µm. (C) The fluorescence intensity for acidic lysosome was normalized to the number of nuclei. ns, not significant. (D) Representative images of acidic lysosomes in controls (41 PDL), MG132-treated, and AA co-treated cells on day 3. Cells were stained with AcidiFluor ORANGE (red) and Hoechst 33342 (blue). Scale bar = 50 µm. (E) Immunoblot analyses of HSP70, LC3, and p62/SQSTM1 for controls, MG132-treated (-), and AA co-treated (+) cells. Intensities of immunoblot bands of LC3-II (F) and HSP70 (G) were normalized to those of LC3-I or GAPDH, respectively. Relative values are shown as the mean (SD) (*n* = 4).

### Antimycin A attenuates UPR^mt^ in MG132-treated cells

MG132 induces genes involved in proteotoxic stress responses, known as unfolded protein response (UPR). To investigate whether AA co-treatment affected the UPR^mt^ in MG132-treated cells, we performed qPCR analyses of *HSPD1* and *HSPE1*, two hallmark genes of the canonical UPR^mt^ pathway [19]. Both genes were highly upregulated as early as 6 h after treatment with MG132. Co-treatment with AA significantly suppressed the induction of both genes at 6 and 12 h (Fig. 4A and 4B). We also examined SOD2 induction, which is known as the SIRT3/FOXO3A axis response to UPR^mt^ [20, 21]. SOD2 protein and mRNA levels gradually increased during MG132 treatment. This induction was also suppressed by AA co-treatment on days 3 and 5 (Fig. 4C-E). Nuclear respiratory factor 1 (NRF1) protein, known as the UPR^IMS^-ERα axis response [21] and which regulates proteasome levels [22] and mitochondrial respiration [23], was slightly induced as early as 3 and 6 h after MG132 treatment, but no notable difference was observed between MG132-treated and AA co-treated cells (Fig. 4F).

**FIGURE 4.**
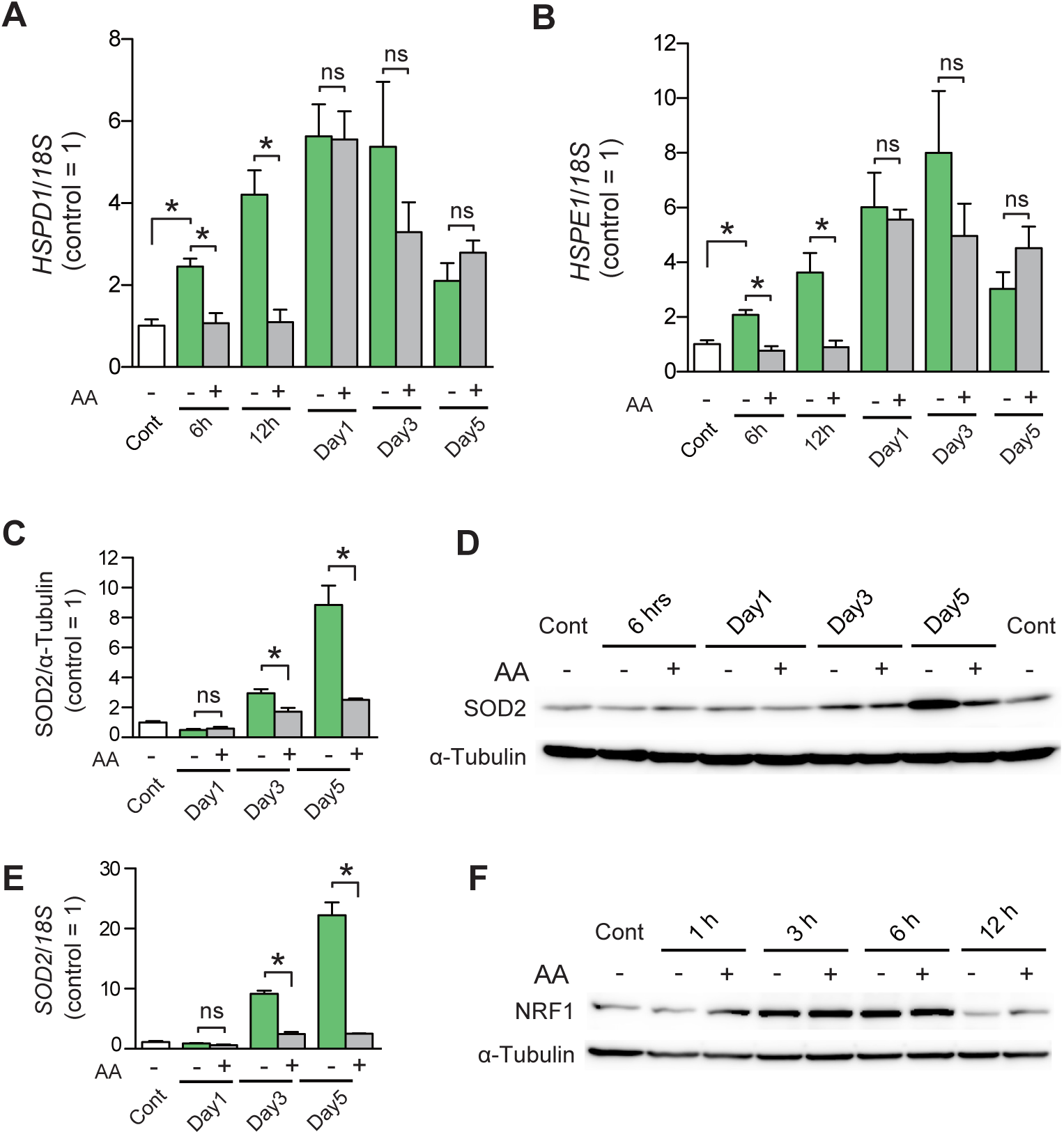
Antimycin A suppresses canonical and late UPR^mt^ but not UPR^IMS^ induced by prolonged inhibition of proteostasis AA co-treatment significantly suppressed the UPR^mt^ elicited by MG132 treatment. *HSPD1* (A) and *HSPE1* (B) mRNA levels in MG132-treated (-) and AA co-treated (+) cells were evaluated by quantitative RT-PCR of four individual cDNAs. The values obtained for *HSPD1* and *HSPE1* were normalized to those of *18S* rRNA. Values are shown as the mean (SD) (*n* = 4). The asterisk (*) indicates *p* < 0.05, as determined by the Mann-Whitney *U* test. ns, not significant. Protein (C and D) and mRNA (E) levels of SOD2 were evaluated by immunoblotting and quantitative RT-PCR analyses of three individually prepared samples. The intensity of SOD2 protein bands was normalized to that of α-tubulin. The value obtained for *SOD2* was normalized to that of *18S* rRNA. Values are presented as the mean (SD). The asterisk (*) indicates *p* < 0.05 by an unpaired t-test. ns, not significant. (F) Immunoblot analysis of NRF1 in MG132-treated (-) vs. AA co-treated (+) cells.

### Antimycin A recovered the temporal depletion of SOD2 in the mitochondrial fraction on day 1 of MG132 treatment

Mitochondrial protein import is inhibited by various oxidative and proteotoxic stresses, including UPR^mt^ and integrated stress response (ISR) [21, 24]. Thus, we investigated the protein levels of two mitochondrial antioxidant enzymes, SOD2 and glutathione peroxidase 4 (GPx4), in the isolated mitochondrial fraction of MG132-treated cells (detailed method is described in Figs. S2A-C). On day 1, SOD2 was significantly depleted in the mitochondrial fraction (Mt) (Figs. 5A and 5B) in cells solely treated with MG132, although the overall SOD2 level (whole) on day 1 was unchanged compared with control cells (Figs. 5A and 5C). On the other hand, GPx4 protein levels were significantly declined not only in Mt but also in the whole fraction on day 1 (Figs. 5A-C). *GPx4* mRNA level was also reduced on day 1 in MG132-treated cells (Fig. 5D). We also observed that SOD2 partially accumulated in the extra-mitochondrial (COX4-negative) regions in MG132-treated cells on day 1, but not in control cells (Fig. 5E). In AA co-treated cells, SOD2 depletion in the mitochondrial fraction on day 1 was restored (Fig. 5F), whereas the whole SOD2 protein level was unchanged by AA co-treatment (Fig. 5G), suggesting that AA recovered the mitochondrial import of SOD2.

**FIGURE 5.**
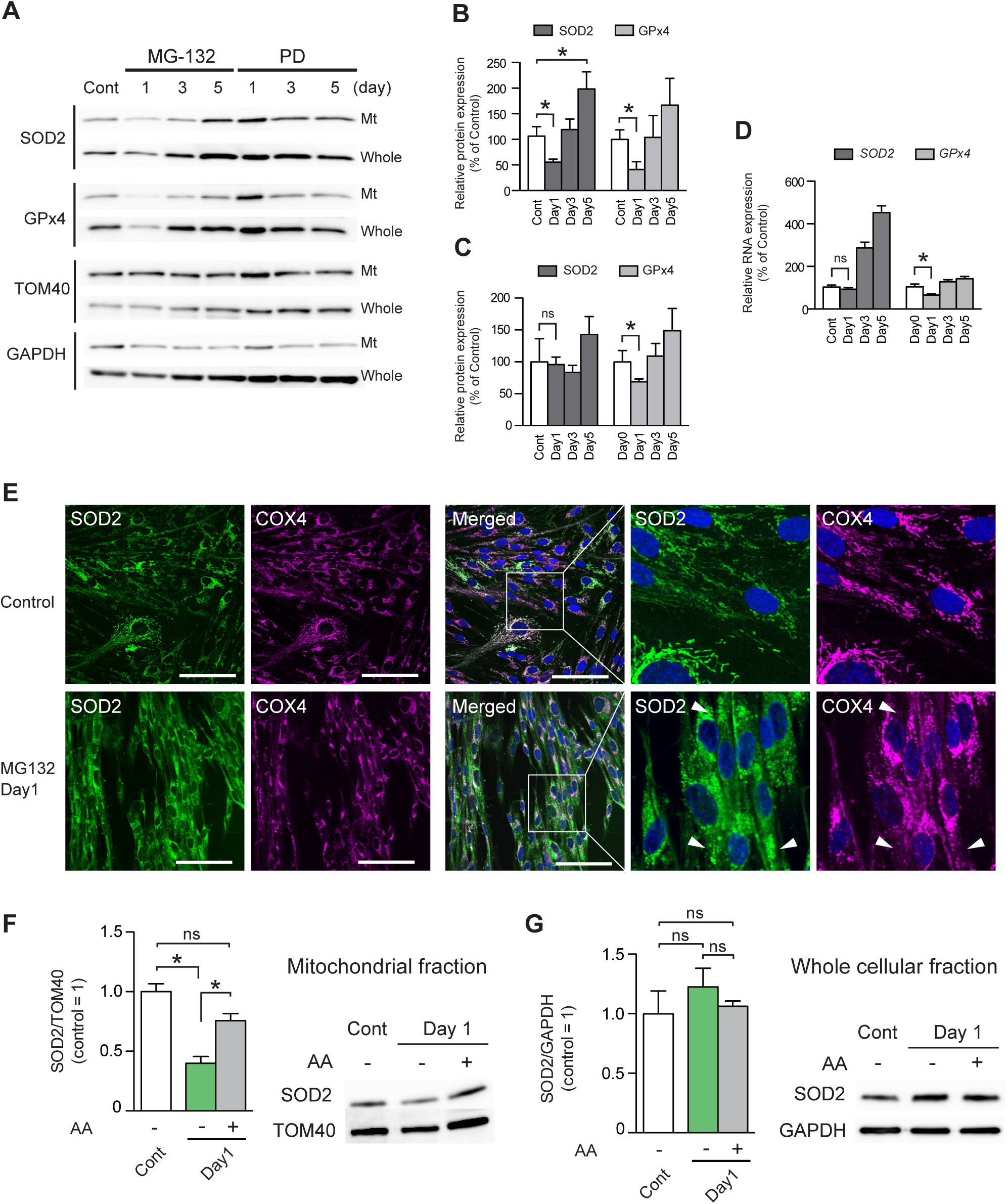
Antimycin A restores temporal depletion of SOD2 in mitochondrial fraction on day 1 of MG132 treatment Depletion of the mitochondrial antioxidant enzymes SOD2 and GPx4 in the mitochondrial fraction of MG132-treated MRC-5 cells in the early period. (A) Immunoblot analyses of MG132-treated cells for SOD2, GPx4, TOM40 (mitochondrial loading control), and GAPDH (cytosolic loading control). Mt, crude mitochondrial fraction; Whole, whole cellular extract; Cont, control cells. Five (mitochondrial fraction) or 15 (whole cellular extract) µg of protein was loaded onto each lane. (B) Quantitative analysis of the mitochondrial SOD2 and GPx4 protein levels in MG132-treated cells. The intensities of the SOD2 and GPx4 bands were normalized to those of TOM40. Values are shown as the mean (SD) (*n* = 4). The asterisk (*) indicates *p* < 0.05. (C) Protein levels of whole-cell SOD2 and GPx4. The intensities of the SOD2 and GPx4 bands were normalized to those of GAPDH. Values are shown as the mean (SD) (n = 4). ns, not significant. (D) The expression levels of *SOD2* and *GPx4* mRNA were evaluated by quantitative RT-PCR of four individual cDNAs prepared from total RNA. (E) Representative images of control and MG132-treated MRC-5 cells immunostained for SOD2 (green), COX4 (purple), and Hoechst 33342 (blue) on day 1. Arrowheads indicate extra-mitochondrially localized SOD2. Scale bar = 100 µm. (F) AA co-treatment restored the mitochondrial depletion of SOD2 protein in MG132-treated cells on day 1. The intensities of SOD2 bands were normalized to those of TOM40. Values are shown as the mean (SD) (*n* = 4). (G) AA co-treatment did not alter whole-cell SOD2 protein expression levels.

### Antimycin A promotes mitochondrial biogenesis and preserves mitochondrial function

We next studied the effect of AA co-treatment on mitochondrial biogenesis and function in MG132-treated cells. Mitochondrial membrane potential (Δψm) was significantly decreased in MG132-treated cells on day 3, as we previously reported [18], and was also reduced in AA co-treated cells (Fig. 6A and 6B). Notably a low Δψm value, a hallmark of mitochondrial dysfunction, in MG132-treated cells suggests the accumulation of defective mitochondria and promotion of mitochondrial clearance. Thus, we investigated mitophagy in MG132- and AA co-treated cells. Mitophagy was remarkably induced in both MG132-treated and AA co-treated cells; however, mitophagy level in AA co-treated cells was significantly lower than that of MG132-treated cells (Fig. 6C and 6D). These results imply enhanced removal of damaged mitochondria predominantly in MG132-treated cells. We previously reported an increase in the mitochondrial DNA (mtDNA) copy number in MG132- and BAFA1-treated cells [18]. In cells co-treated with AA or rotenone, but not CCCP, mtDNA copy number was significantly higher than that in MG132-treated cells (Fig. 6E). These results suggest that AA co-treatment more strongly shifted the balance between mitochondrial clearance and biogenesis, in favor of increased biogenesis. In fact, the protein levels of NRF1 and TFAM (also known as mtTFA), which are critically involved in mitochondrial biogenesis [23], were notably increased in AA co-treated cells (Figs. 6F-6H). Furthermore, DRP1, a key mitochondrial division-promoting factor, was also upregulated in AA co-treated cells (Fig. 6F and 6I), suggesting that damaged parts of the mitochondria were divided and eliminated by mitophagy. On the other hand, BNIP3 and Nix, major regulators of receptor-mediated mitophagy and cell death [25, 26], were significantly upregulated in MG132-treated cells on days 1 and 3, while the expression levels of these proteins in AA co-treated cells were similar to those in control cells (Fig. 6F, 6J, and 6K). Parkin protein expression was largely unchanged in all samples (Fig. 6F), indicating that damaged mitochondria were eliminated through the Parkin-independent pathway in MG132-treated cells. Mitochondrial morphology is closely associated with various mitochondrial and cellular functions [27]. We further investigated mitochondrial morphology to assess whether the mitochondrial dynamics observed in MG132- and AA co-treated cells were associated with mitochondrial structure. For this purpose, we stained cells with MitoTracker Green FM and analyzed fluorescent images (Fig. 6M) using MiNA, the mitochondrial network morphology analysis toolset [25]. The results revealed significant increases in the mitochondrial footprint in MG132-treated and AA co-treated cells (Fig. 6N), suggesting an increase in mitochondrial mass in AA co-treated and MG132-treated cells. This finding supports the prediction that mitochondrial biogenesis is enhanced by AA co-treatment. The mean branch length of AA co-treated cells was also longer than that of control and MG132-treated cells (Fig. 6O). The mean network branch of AA co-treated cells was significantly higher than that of the other two groups, whereas that of MG132-treated cells was comparable to that of control cells (Fig. 6P). This possibly indicates that AA increased the branched network structure of mitochondria rather than individual structures such as rods and large/round structures that were predominantly observed in MG132-treated cells (Fig. 6M).

**FIGURE 6.**
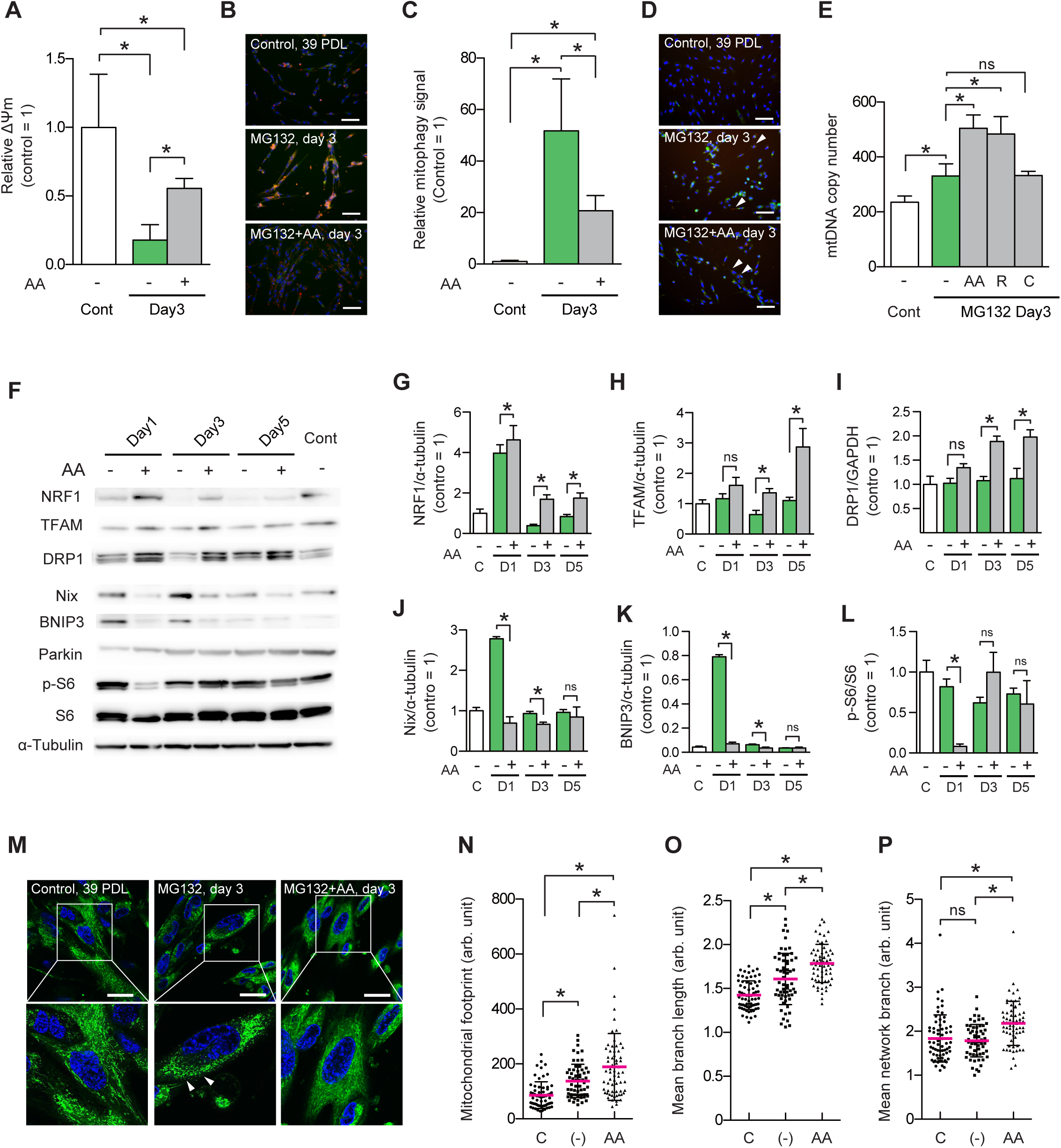
Antimycin A suppresses mitophagic flux induced by prolonged inhibition of proteostasis and enhances mitochondrial biogenesis (A) Relative mitochondrial membrane potential (ΔΨm) was evaluated using JC-1 MitoMP Detection Kit. Normalized fluorescence intensity for JC-1 dye (red/green) was quantified using FIJI/ImageJ2. JC-1 dye emits red fluorescence upon potential-dependent accumulation in mitochondria. Relative values are shown as the mean (SD) from the measurements of the four different images. The asterisk (*) indicates *p* < 0.05, using one-way ANOVA with Tukey’s multiple comparison test. (B) Representative images of ΔΨm signal in control (39 PDL), MG132-treated, and AA co-treated cells on day 3. Cells were stained with JC-1 dye (red and green) and Hoechst 33342 (blue). Scale bar = 50 µm. (C) Relative mitophagy signal was evaluated using Mitophagy Detection Kit. The fluorescence intensities for Mtphagy dye and Hoechst 33342 were quantified using FIJI/ImageJ2. Red fluorescence intensity was normalized to the number of nuclei. Relative values are shown as the mean (SD) from the measurements of the six different images. The asterisk (*) indicates *p* < 0.05. (D) Representative images of mitophagy signals in control (39 PDL), MG132-treated, and AA co-treated cells on day 3. Cells were stained with Mtphagy dye (red), Lyso dye (green), and Hoechst 33342 (blue). Arrowheads indicate mitophagy signals. Scale bar = 50 µm. (E) The copy number of mitochondrial DNA (mtDNA) of control, MG132-treated, and co-treated cells with an ETC inhibitor was measured by qPCR using mtDNA-specific primers and nuclear DNA-specific primers as a reference. Values are the mean (SD) (*n* = 4). The asterisk (*) indicates *p* < 0.05. (F) Immunoblot analyses of proteins related with mitochondrial functions, for controls, MG132-treated (-), and AA co-treated (+) cells. Intensities of immunoblot bands of NRF1 (G), TFAM (H), DRP1 (I), Nix (J), BNIP3 (K), and phosphorylated ribosomal protein S6 (p-S6) (L) were normalized, respectively. Relative values are shown as the mean (SD) (*n* = 4). (M) Representative images of MitoTracker Green-stained mitochondrial morphology in control, MG132-treated, and AA co-treated cells on day 3. Arrowheads indicate large punctate structures of mitochondria. Scale bar = 30 µm. Statistical analyses for mitochondrial footprint (N), mean branch length (the mean length of all the lines used to represent the mitochondrial structures) (O), and mean branch length (the mean length of all the lines used to represent the mitochondrial structures) (P) in control, MG132-treated, and AA co-treated cells on day 3. Scatter dot plots show mean (horizontal red line) with SD. Mitochondrial network analysis (MiNA) was performed on 50-60 cells. The asterisk (*) indicates *p* < 0.05, using one-way ANOVA with Tukey’s multiple comparison test.

## Discussion

Mitochondria are the bioenergetic center and the main source of intracellular reactive oxygen species (ROS). As we have previously shown, mitochondria, despite being intact or dysfunctional, accumulate during SIPS progression caused by prolonged disturbance of proteostasis, along with the production of excess ROS [18]. However, the involvement of mitochondrial respiration, one of the most pivotal mitochondrial functions, in the SIPS is unknown. Here, we revealed that inhibition of complex III in the ETC by antimycin A resulted in notable attenuation of SIPS induced by prolonged treatment of human fibroblasts with MG132 or BAFA1. The levels of intracellular ROS and protein aggregates were significantly suppressed by AA co-treatment, implying the crucial involvement of mitochondrial respiration in intracellular ROS production and protein quality control. When cells were co-treated with vitamin E to scavenge ROS in our SIPS model, intracellular H_2_O_2_ and protein aggregate levels were concomitantly suppressed [18]. Therefore, it is plausible that the reduction in protein damage caused by ROS can contribute to the reduced protein aggregate levels in AA co-treated cells.

MG132 treatment triggers unfolded protein response in endoplasmic reticulum (UPR^er^)[28] and mitochondria (UPR^mt^)(Fig. 4). Interestingly, we found that the upregulation of two typical target genes, *HSPD1* and *HSPE1*, of canonical UPR^mt^ by MG132 was significantly suppressed at very early periods of AA co-treatment, but induced later, up to comparable levels of single treatment with MG132, suggesting that delayed induction of UPR^mt^ is relevant to SIPS attenuation. NRF1 expression was not altered by AA treatment.

Rawat *et al*. reported that nuclear-encoded mitochondrial proteins such as respiratory chain complex subunits form cytosolic aggregates as an early event in MG132-treated cells [29]. We observed marked depletion of the antioxidant enzymes SOD2 and GPx4 in the mitochondrial fraction (Figs. 5A and 5B) and notable cytosolic localization of SOD2 in MG132-reated cells on day 1 (Fig. 5E). Extra-mitochondrial SOD2 also showed partial colocalization with aggresome-like inclusions and lysosomes (data not shown) in MG132-treated cells on day 1, implying that cytosolic SOD2 was insufficiently translocated into the mitochondria as an early event of disturbance of proteostasis. On the other hand, GPx4 protein levels in both mitochondrial and whole fractions, along with *GPx4* mRNA levels, were significantly reduced on day 1. Thus, GPx4 depletion in the mitochondrial and whole fractions possibly arose from the reduction in GPx4 transcripts. In AA co-treated cells, mitochondrial depletion of SOD2 on day 1 was restored (Fig. 5F). Considering the suppression of intracellular protein aggregates by co-treatment with AA (Fig. 3A and 3B), it is likely that the overall protein integrity was preserved in AA co-treated cells, resulting in proper SOD2 localization. However, the detailed mechanism of mitochondrial import in MG132- and AA co-treated cells remains unclear.

MG132 treatment highly induced mitophagy on day 3, which was possibly triggered by the accumulation of defective mitochondria with low ΔΨm (Fig. 6A-D). The remarkable upregulation of mitophagy in MG132-treated cells might be mediated by the NIX/BNIP3-dependent pathway but not by the Parkin-dependent pathway (Fig. 6F). Although the mitophagy level of AA co-treated cells on day 3 was also higher than that of control cells, it was significantly lower than that of MG132-treated cells (Fig. 6C and 6D). Furthermore, co-treatment with AA appeared to enhance mitochondrial biogenesis through NRF1 upregulation (Fig. 6F and 6G), which resulted in an increase in TFAM protein levels (Fig. 6F and 6H). DRP1 protein levels (Fig. 6I) and mtDNA copy number (Fig. 6E) in AA co-treated cells notably increased on day 3 when compared with those of MG132-treated cells, implying that mitochondria in AA co-treated cells sustain their function by properly regulating the clearance of defective mitochondria by mitophagy and regeneration of functional mitochondria through mitochondrial biogenesis. We previously showed that the levels of phosphorylated S6 (p-S6), the best marker of mTOR activity, were largely unchanged during MG132 treatment [18]. Upon co-treatment with AA, the p-S6/S6 ratio significantly decreased on day 1 (Fig. 6F and 6L), implying that AA rapidly arrested cell growth and switched to the catabolic state.

Thereafter, p-S6 returned to a level comparable to that of the control on days 3 and 5, supporting the promotion of mitochondrial biogenesis by shifting the cellular metabolism to the anabolic state. Numerous studies over the past two decades have revealed a close relationship between mitochondrial respiration and longevity [30]. Pharmacological inhibition or genetic ablation of components in mitochondrial ETC complexes results in extended lifespans in several organisms. In contrast, inhibition of ETC induces mitochondrial stress and causes cellular senescence in many cases [31]. Sustained inhibition of ETC complex III by AA arrested cell proliferation and induced premature senescence [32]. In our study, temporal inhibition of ETC complex III by AA in the proteostasis-disturbed model resulted in significant attenuation of SIPS (Figs. 1D-K). Although we revealed that AA co-treatment with MG132 reduced the rate of UPR^mt^ induction (Figs. 4A-F) and enhanced mitochondrial biogenesis (Figs. 6E-N), the detailed molecular mechanisms of retrograde signaling from inhibition of mitochondrial respiration by AA to changes in nuclear gene regulation could not be unveiled (arrow with “?” in the graphical abstract). In our model, the levels of mitochondrial superoxide and intracellular H_2_O_2_ were suppressed by AA co-treatment (Figs. 2B-C), but this did not appear to be caused by the upregulation of antioxidant enzymes such as SOD2 (Fig. 4C and 5A-C). Therefore, one explanation is that AA could prevent excess ROS production by an unknown mechanism, which might be sufficient for the attenuation of SIPS by disturbed proteostasis. Alternatively, inhibition of ETC complex III might hamper the oxidation of ubiquinone (CoQ10), resulting in the accumulation of the reduced form of CoQ10, which functions not only as an electron-transferring molecule for ETC complexes I-III, but also as an antioxidant molecule and exerts an anti-senescence effect in mice [33] and cultured cells [34]. However, our preliminary experiments, in which MRC-5 cells were co-treated with MG132 and the reduced form of CoQ10 (10 µM) for five consecutive days, showed no differences in SA-β-gal activity between MG132-treated and CoQ10-co-treated cells (Fig. S3). Therefore, studying other mitochondrial metabolites, such as NAD^+^ and α-ketoglutarate, might uncover the protective mechanism of AA co-treatment in the proteostasis-disturbed cellular senescence model.

In conclusion, we have shown that temporal inhibition of mitochondrial respiration by ETC inhibitors significantly attenuated SIPS induced by impaired proteostasis, in which intracellular ROS levels, acute UPR^mt^ responses, and accumulation of protein aggregates were remarkably suppressed. Furthermore, AA co-treatment recovered the temporal depletion of SOD2 in the mitochondrial fraction, suppressed the induction of mitophagy, and enhanced mitochondrial biogenesis. These findings provide strong evidence that mitochondrial respiration is closely associated with the progression of premature senescence caused by impaired proteostasis.

### Experimental procedures

#### Cell culture

MRC-5 cells (30 PDL) were obtained from the JCRB Cell Bank. The cells were maintained in EMEM (Fujifilm Wako Pure Chemical Corp, Osaka, Japan) supplemented with 10% fetal bovine serum (FBS)(Biological Industries, Cromwell, CT) and penicillin-streptomycin (Nacalai Tesque, Kyoto, Japan) at 37°C in a humidified atmosphere of 5% CO_2_. The cells were passaged every 2–3 days.

#### MG132 and BAFA1 treatment

For MG132 or BAFA1 treatment, we plated 1 × 10^4^ cells per well in a 96-well clear-bottom black plate, allowed them to grow for 16–24 h, and then changed the culture medium to EMEM containing MG132 (ChemScene, Monmouth Junction, NJ) or BAFA1 (Sigma-Aldrich). As a pilot experiment, we tested the viability of cells treated with MG132 (0.2–0.4 µM) or BAFA1 (8–15 nM) at several different concentrations to determine the appropriate concentration of each drug. On day 5 of drug treatment, the culture medium was replaced with regular EMEM containing no drug. For co-treatment with ETC inhibitors, the cells were treated with 1 µM antimycin A (Sigma-Aldrich), 1 µM rotenone (Sigma-Aldrich), or 1 µM CCCP (Sigma-Aldrich).

#### Immunoblotting and antibodies

Cells cultured in 6-cm dishes were trypsinized and harvested. The cell pellets were lysed in RIPA buffer (20 mM Tris-HCl, pH7.4, 100 mM NaCl, 0.5% Nonidet P-40, 0.05% SDS, 1 mM EDTA, and protease inhibitors). Protein concentrations were determined using Quick Start Bradford dye (Bio-Rad). Whole cell extracts (10–20 µg) were mixed with 5×SDS sample buffer containing 2-mercaptoethanol, heated at 95°C for 3 min, and loaded onto an SDS-polyacrylamide gel (SDS-PAGE). Proteins were transferred to a PVDF membrane and incubated with 1% Western blocking reagent (Sigma-Aldrich) at 25°C for 1 h. The membrane was then incubated overnight with antibodies diluted in 0.5% Western blocking reagent. The antibodies used in this study are summarized in the Supplemental file. The chemiluminescence signal was visualized with ECL prime (Cytiva) or ImmunoStar LD (Fujifilm), and detected with LAS4000 (GE Healthcare) or ImageQuant800 (Cytiva). The intensity of the obtained protein band was quantified using FIJI/ImageJ2.

#### SA-β-gal staining and senescence assays

SA-β-gal staining was performed as previously described [35]. Quantitation of SA-β-gal activity was performed using the Cellular Senescence Plate Assay Kit - SPiDER-βGal, a fluorogenic substrate for β-galactosidase (Dojindo, Kumamoto, Japan), following the manufacturer’s instructions.

#### Immunofluorescence

For immunofluorescence analyses, cells were fixed in 100% methanol, incubated with Blocking One Histo (Nacalai Tesque, Kyoto, Japan) for 1 h, and then incubated with diluted primary antibody overnight. The antibodies used in this study are summarized in the Supplemental file. After several TBS-T washes, cells were incubated with a secondary antibody and stained with Hoechst 33342 (Dojindo, Kumamoto, Japan). Cells were observed under an all-in-one fluorescence microscope BZ-9000 (Keyence, Osaka, Japan), and fluorescence images were quantified using FIJI/ImageJ2.

#### Measurement of intracellular H_2_O_2_

Cells plated in a 24-well plate were incubated with 1 µM HYDROP (Goryo Chemical, Sapporo, Japan) with Hoechst 33342 diluted in EMEM containing FBS for 30 min at 37°C, and then washed once with 500 µL Hank’s balanced salt solution (HBSS). Fluorescence (Ex/Em = 485/535 nm) was observed under BZ-9000 (Keyence, Osaka, Japan), and fluorescence images were quantified using FIJI/ImageJ2. Normalized values were obtained by dividing the fluorescence intensities of HYDROP by the number of nuclei stained with Hoechst 33342.

#### Staining of protein aggregation

Protein aggregation was visualized using ProteoStat Aggresome Detection Reagent (Enzo Life Science, Farmingdale, NY) according to the manufacturer’s protocol. Cells plated in 48-well plate were observed under BZ-9000 (Keyence, Osaka, Japan), and fluorescence images were quantified using FIJI/ImageJ2.

#### Live cell microscopy for mitochondrial analyses

To detect ΔΨm, live cells were incubated with 2 µM JC-1 MitoMP detection reagent (Dojindo, Kumamoto, Japan) with Hoechst 33342 diluted in EMEM containing FBS for 30 min at 37°C, and then washed once with 100 µL HBSS. To detect mitochondrial ROS, cells were incubated with 5 µM MitoSOX Red mitochondrial superoxide indicator (Thermo Fisher Scientific) in HBSS for 10 min at 37°C, washed with HBSS three times, and cultured in regular medium for 6 h. Cells stained with Hoechst 33342 were washed once with 100 µL HBSS, and observed under BZ-9000 (Keyence, Osaka, Japan). Fluorescence images were analyzed and quantified using FIJI/ImageJ2. Mitophagy levels were examined using Mitophagy detection kit (Dojindo, Kumamoto, Japan). Cells were stained with 100 nM Mitophagy dye solution in HBSS for 30 min at first, washed twice with HBSS, and then stained with 1 µM Lyso dye solution and Hoechst 33342 in HBSS for 30 min at 37°C, and observed under BZ-9000. Only mitophagy fluorescence superimposed on the lysosomal signal was quantified using FIJI/ImageJ2.

#### Intracellular ATP assay

Intracellular ATP levels were measured using “Cell” ATP Assay reagent (Toyo B-Net, Tokyo, Japan). The ATP level was normalized to the DNA content in the cell lysates, which was quantified by SYBR Gold staining (Thermo Fisher Scientific).

### Copy number of mitochondrial DNA

Total DNA was isolated from MRC-5 cells using DNeasy Blood & Tissue kit (QIAGEN). To quantify the copy number of mitochondrial DNA, we performed qPCR analyses using Human Mitochondrial DNA Monitoring Primer set (TAKARA BIO, Shiga, Japan) and Thermal Cycler

Dice Real Time System III (TAKARA BIO, Shiga, Japan).

### Preparation of mitochondrial fraction

The procedure for isolation of the mitochondrial fraction from cultured cells was as described by Clayton *et al.* [36] with slight modifications. The buffer volume, number of strokes, homogenizer, and pestle were optimized for 0.5–1.0 × 10^6^ cells. Frozen cell pellets were resuspended in a hypotonic buffer and homogenized using a disposable plastic pestle (As One Corp., Osaka, Japan) with matched Safe-Lock tubes (Eppendorf). The detailed procedure is summarized in Figs. S2A-C.

### Quantitative PCR

Total RNA was isolated from MRC-5 cells using ISOGEN reagent (Nippon gene, Tokyo, Japan) and reverse-transcribed using ReverTra Ace qPCR RT Master Mix with gDNA Remover (TOYOBO, Osaka, Japan). cDNAs of human *SOD2, GPx4,* and *18S* ribosomal RNA were amplified using THUNDERBIRD Next SYBR qPCR Mix (TOYOBO, Osaka, Japan) and Thermal Cycler Dice Real Time System III. Primer sequences are listed in the Supporting Information.

### Statistical analyses

We conducted one-way ANOVA followed by Tukey’s multiple comparison test. The two groups were compared using the non-parametric Mann-Whitney *U* test. Differences between the groups were considered significant at *p* < 0.05. All data were analyzed using the GraphPad Prism 5.0 software (GraphPad Software, San Diego, California USA, www.graphpad.com) and presented as the mean (SD) of the obtained values.

### Mitochondrial morphology analysis

The mitochondrial morphology of MRC-5 cells was analyzed using the Mitochondrial Network Analysis toolset (MiNA) version 3.0.1 available at https://github.com/StuartLab/MiNA run on ImageJ2/FIJI. MRC-5 cells were seeded in µ-Slide 8 well chamber slide (ibidi GmbH, Germany) and treated with MG132 and antimycin A. On day 3, live cells were stained with 0.5 µM MitoTracker Green FM (Thermo Fisher Scientific) with Hoechst 33342 diluted in EMEM containing FBS for 30 min at 37°C, and then washed with HBSS once. The cells were observed under a laser-scanning microscope FV1200 (Olympus, Tokyo, Japan). Images were opened with FIJI/ImageJ2, processed to 8-bit grayscale images, and preprocessed using an unsharp mask filter. A single cell was selected using an ROI manager, and the mitochondrial morphology of each cell was analyzed using the MiNA macro plugin with a Ridge Detection (2D) function to generate output parameters. Measurements were visualized and statistically analyzed using the GraphPad Prism 5.0 software.

## Acknowledgements

The authors are grateful to Santa Cruz Biotechnology, Inc. (Dallas, TX) and Cosmo Bio Co., Ltd. (Tokyo, Japan) for providing the sample antibodies listed in Supporting Information. The authors would like to thank Editage (www.editage.jp) for English language editing, Mr. Hiroki Takenaka for aiding mitochondrial morphological analyses using MiNA, and KANEKA Corporation (Osaka, Japan) for providing oxidized and reduced CoQ10.

## Author contributions

Y. T. and Y. K. designed the research and drafted the paper; Y. T. carried out experiments and analyzed data; I. I., M. H., and M. I. supported the analysis of oxidative stress.

## Funding info

This work was supported by JSPS KAKENHI Grant Number JP21K11605.

## Data availability statement

The data that support the findings of this study are available from the corresponding author upon reasonable request.

## Conflict of interest

The authors have no conflicts of interest to declare in association with this study.

---

**Figure S1.**
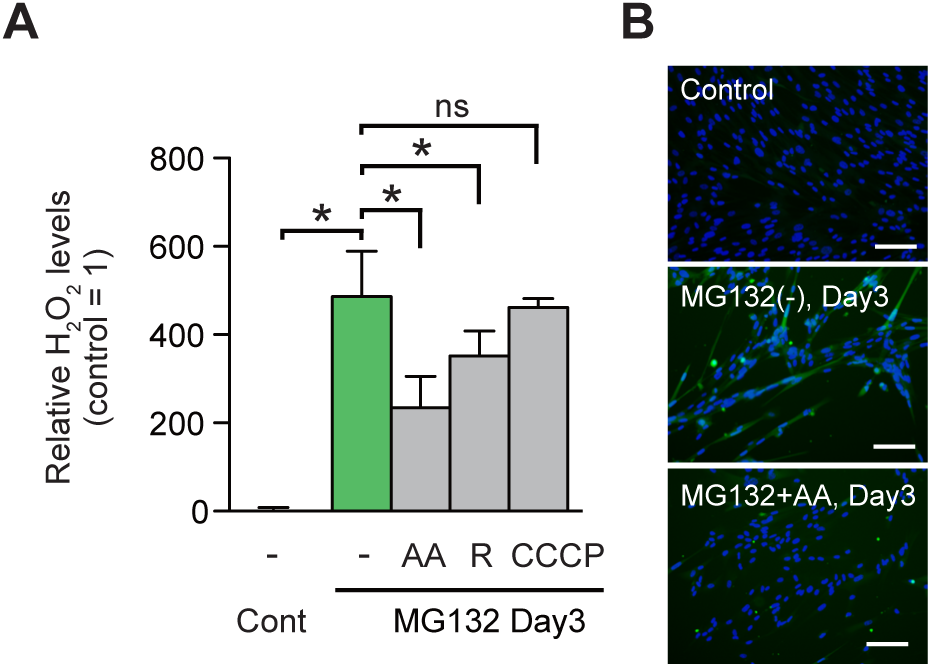
Antimycin A suppresses generation of excess intracellular hydrogen peroxide induced by prolonged inhibition of proteostasis (A) Relative intracellular H_2_O_2_ levels in MG132-treated (-) and antimycin A (AA), rotenone (R), or carbonyl cyanide 3-chlorophenylhydrazone (CCCP) co-treated cells on day 3. The fluorescence intensity for HYDROP was normalized to the number of nuclei. Relative values are shown as the mean (SD) (n = 4). The asterisk (*) indicates p < 0.05 by a Mann-Whitney U test. ns, not significant. (B) Representative images of control, MG132-treated, and AA co-treated cells on day 3 stained with HYDROP (green) and Hoechst 33342 (blue). Scale bar = 50 µm.

**Figure S2.**
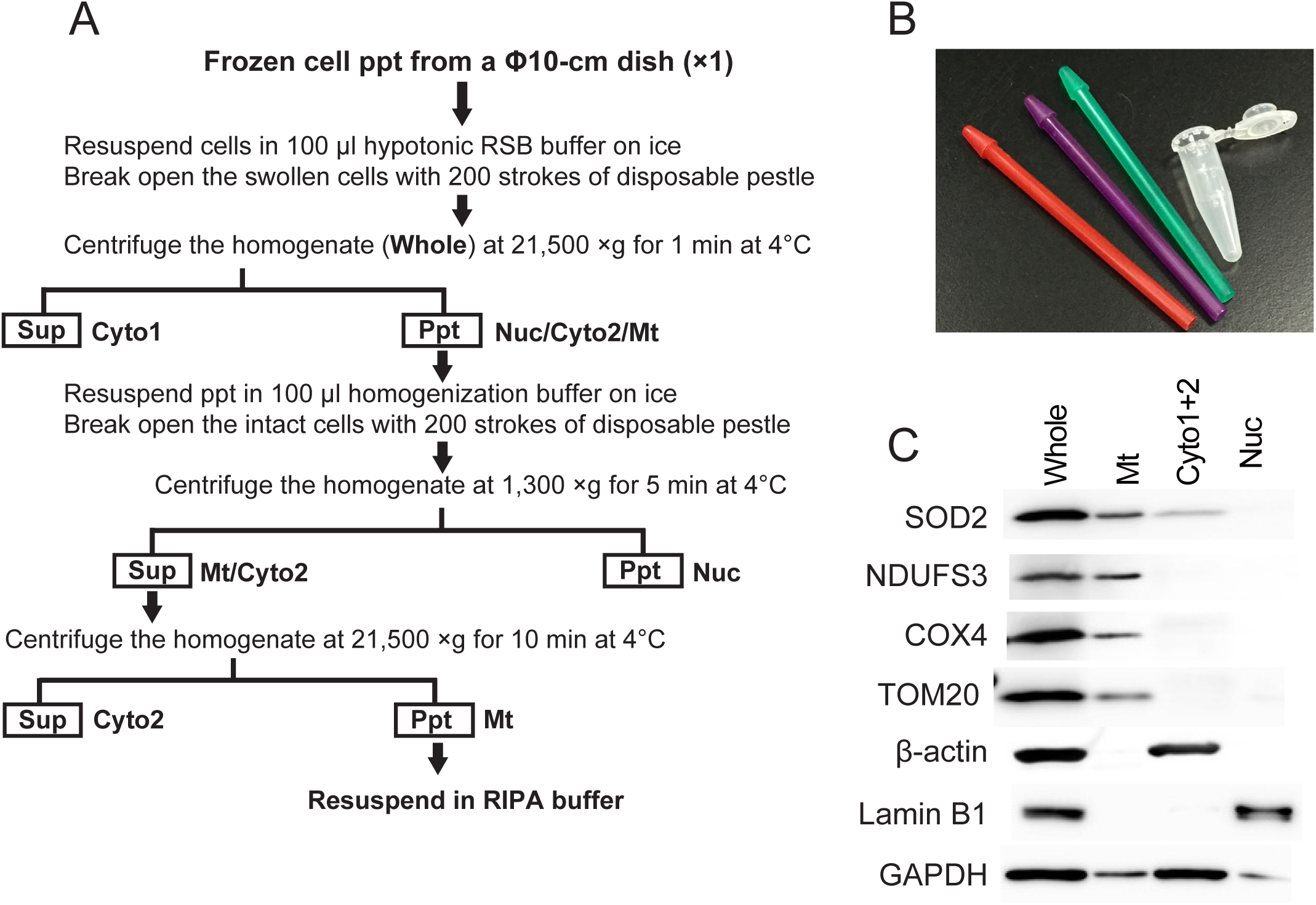
Preparation of crude mitochondrial fraction from MRC-5 cells. (Α) A flow chart for preparation of mitochondrial fraction from frozen cell pellet at a microscale. Cells were homogenized in hypotonic RSB buffer, then unbroken cells were repeatedly homogenized in a homogenization buffer. For the immunoblot analyses, solubilized mitochondrial proteins in RIPA buffer containing 0.5% NP40 (Mt) were applied. Hypotonic RSB buffer, 10 mM Tris-HCl, pH7.4, 10 mM NaCl, 1.5 mM MgCl2, protease inhibiter; Homogenization buffer, 5 mM Tris-HCl, pH7.4, 1.5 mMgCl2, 250 mM sucrose, protease inhibitor; RIPA buffer, 20 mM Tris-HCl, pH7.4, 100 mM NaCl, 0.5% Nonidet P-40, 0.05% SDS, 1 mM EDTA, protease inhibitors; Cyto, cytoplasmic fraction; Nuc, nuclear fraction; Whole, whole cellular extract. (B) Disposable plastic pestles and a matched Safe-Lock tube used in this method. (C) Immunoblot analyses of whole cellular extract (Whole), crude mitochondrial fraction (Mt), and cytoplasmic fraction (Cyto1 and 2) prepared from untreated control MRC-5 cells using antibodies against SOD2, NDUFS3, COX4, TOM20, β-actin, Lamin B1, and GAPDH.

**Figure S3.**
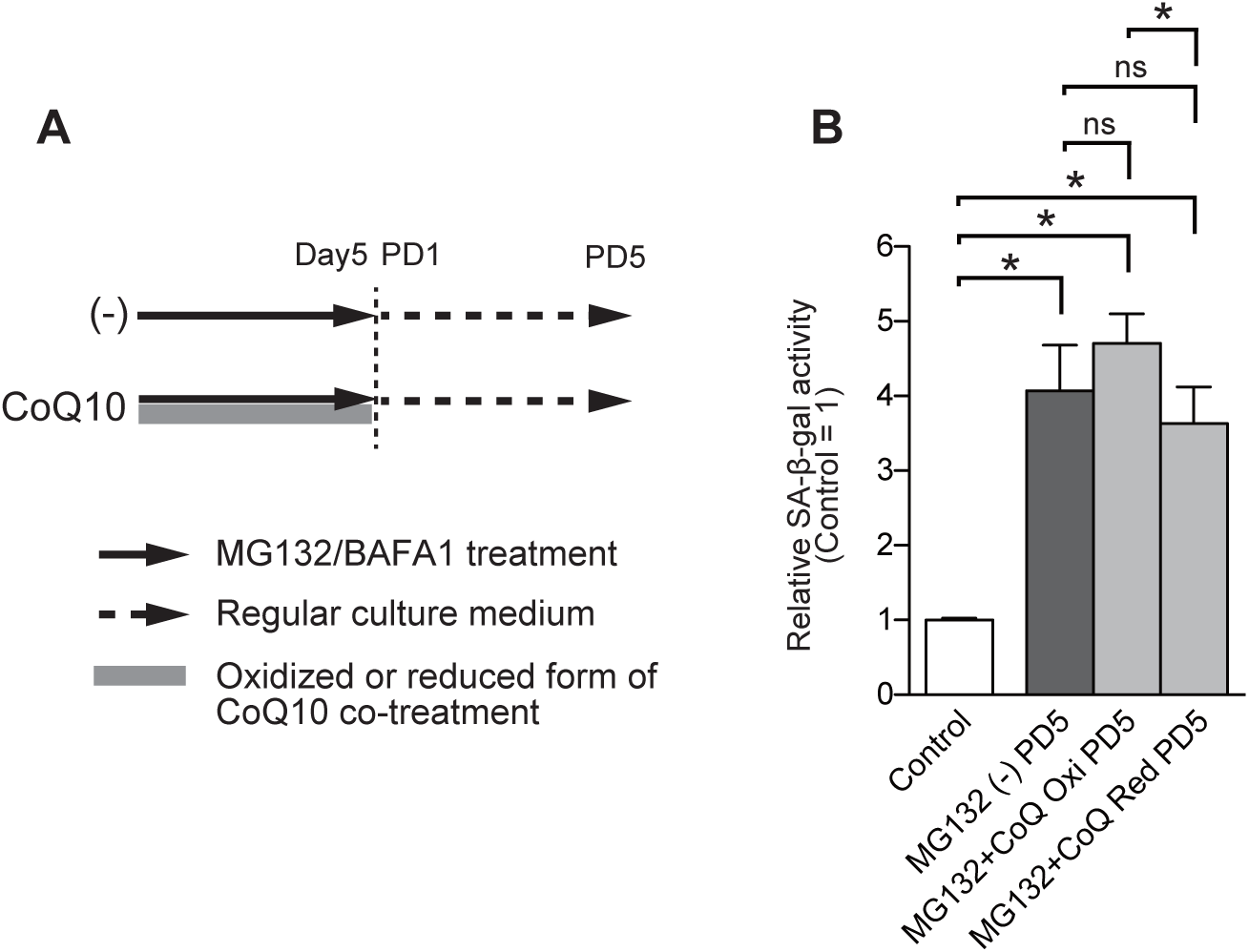
Co-treatment of oxidized (Oxi) or reduced (Red) form of coenzyme Q10 (CoQ) with MG132 does not affect the SA-β-gal activity at PD5 in human MRC-5 cells. Relative SA-β-gal activities in MG132-treated (-) and 10 µM oxidized (Oxi) or reduced (Red) coenzyme Q10 (CoQ) co-treated cells at PD5. The fluorescence intensity for SPiDER-β-Gal was normalized to that of nuclei. Relative values are shown as the mean (SD) (n = 4). The asterisk (*) indicates p < 0.05 by a Mann-Whitney U test. ns, not significant.

## Supporting information

### 1. Antibodies used in this study

#### 1st antibodies

anti-S6 (C-8, sc-74459, Santa Cruz Biotechnology, mouse monoclonal)

anti-p-S6 (50.Ser 235/236, sc-293144, Santa Cruz Biotechnology, mouse monoclonal)

anti-NRF1 (G-5, sc-515360, Santa Cruz Biotechnology, mouse monoclonal)

anti-NRF2 (M200-3, MBL Life Science, mouse monoclonal)

anti-LC3 (PM036, MBL Life Science, rabbit polyclonal)

anti-GAPDH (G8795, Sigma-aldrich, mouse monoclonal)

anti-AKT (#9272S, Cell Signaling Technology, rabbit polyclonal)

anti-p-AKT (S473) (#9271S, Cell Signaling Technology, rabbit polyclonal)

anti-SQSTM1 (p62) (D-3, sc-28359, Santa Cruz Biotechnology, mouse monoclonal)

anti-HSP70 (C92F3A-5, sc-66048, Santa Cruz Biotechnology, mouse monoclonal)

anti-NIX (H-8, sc-166332, Santa Cruz Biotechnology, mouse monoclonal)

anti-BNIP-3 (ANa40, sc-56167, Santa Cruz Biotechnology, mouse monoclonal)

anti-Parkin (PRK8, sc-32282, Santa Cruz Biotechnology, mouse monoclonal)

anti-TFAM (C-9, sc-376672, Santa Cruz Biotechnology, mouse monoclonal)

anti-DRP1 (C-5, sc-271583, Santa Cruz Biotechnology, mouse monoclonal)

#### 2nd antibodies

HRP-conjugated goat anti-mouse IgG (A9044, Sigma-Aldrich) HRP-conjugated anti-rabbit IgG (NA934, GE healthcare)

DyLight 488-conjugated anti-mouse IgG (#35503, Thermo Fisher, mouse monoclonal)

### 2. Quantitative PCR (qPCR)

#### qPCR primers

For human SOD2

Hs_SOD2_RT-UP1, 5’-AACGGGGACACTTACAAATTGCT-3’ Hs_SOD2_RT-LP1, 5’-CCCAGTTGATTACATTCCAAATAGCTT-3’

For human GPx4

Hs_GPx4_RT-UP1, 5’-CGCGGGCTACAACGTCAAATTCGAT-3’ Hs_GPx4_RT-LP1, 5’-CCACGCAGCCGTTCTTGTCGAT-3’

For 18S ribosomal RNA

LEM-m18S-F, 5’-CGGCTACCACATCCAAGGAA-3’ LEM-m18S-R, 5’-GCTGGAATTACCGCGGCT-3’

## Notes

### Competing Interest Statement

The authors have declared no competing interest.

